# Transcriptome and metabolome analyses reveal pathways associated with fruit color in plum (*Prunus salicina Lindl*.)

**DOI:** 10.1101/2021.07.23.453563

**Authors:** Lei Chen, XueSong Wang, Long Cui, YanBo Zhang

## Abstract

**Background:** In order to reveal the mechanism of fruit color changes in plum, two common plum cultivars *Changli84* (Ch84, red fruit) and *Dahuangganhe* (D, yellow fruit) in Northeast China were selected as plant materials. Transcriptome sequencing and metabonomic analyzing were performed at three different developmental stages: young fruit stage, colour-change stage, and maturation stage.

**Results:** “Flavonoid biosynthesis” was significantly enriched in the KEGG analysis. Some DEGs in “Flavonoid biosynthesis” pathway had an opposite trend between the two cultivars, such as *CHS, DFR* and *FLS*. Also, transcriptional control of MBW (MYB–bHLH–WD) protein complexes showed a close relationship with plum fruit color, especially the expression of *MYBs* and *bHLHs*. In the current study, procyanidin B1 and B2 had the highest level at young fruit stage in Ch84 and the content of procyanidin B2 decreased sharply at the color change stage. Conversely, the content of cyanidin increased with the growth of fruit and reached the peak at the maturation stage.

**Conclusion:** The content of procyanidin B1 and B2 in plums at young fruit stage might be the leading factors of the matured fruit color. At the maturation stage, the cyanidin produced by procyanidins keeps the color of the fruit red. Correspondingly, genes in “flavonoid biosynthesis” pathway play critical roles in regulating the accumulation of anthocyanin in plum.

## Introduction

Plums (Prunus salicina Lindl.) is a kind of favorite fruit product by consumers for their delicious tastes. Plum have a wide variety of uses and consumers typically prefer to eat fresh plums for their characteristic taste and rich nutrient substance [1]. There are many varieties of plums, which have different characteristics, such as maturity period, different taste, different fruit color and so on. In this study, two plum cultivars with different fruit color were selected to study the molecular mechanism that related to fruit color formation.

Concentration of anthocyanins determines the color of fruits [2, 3]. Anthocyanins are responsible for the colors of numerous flowers, fruits, vegetables and even cereals [4], and can strongly contribute to food quality and appeal to consumers. Anthocyanins are the most abundant flavonoid compounds, which possess antioxidant and anti-inflammatory properties [5, 6]. Anthocyanin synthesis is regulated by many external environmental factors, such as nutrient depletion, drought, pathogen infection, temperature, light and genetic factors [7]. Different factors could cause content changes in anthocyanin abundance via different pathways. Genetic factors that related to anthocyanin accumulation can be divided into two categories. One category is the biosynthetic genes that encode enzymes required for anthocyanin biosynthesis, such as follows: chalcone synthase (*CHS*), chalcone isomerase (*CHI*), flavanone-3-hydroxylase (*F3H*), dihydroflavonol 4-reductase (*DFR*), anthocyanidin synthase (*ANS*) and UDP glucose flavonoid 3-O-glucosyltransferase (*UF3GT*) [8]. Another category is regulatory genes that influence the intensity and pattern of anthocyanin biosynthetic genes, including two major classes of transcription factors: the myeloblastosis (MYB) and basic helix–loop–helix (bHLH) families, which, along with WD-repeat protein, form a MYB–bHLH–WDR transcription complex to regulate anthocyanin synthesis [9-11]. Some of these structural and regulatory genes had been identified and cloned in model plants such as Arabidopsis, maize (Zea mays), etc. [12-15]. It has been reported that anthocyanins are closely related to peel color, and pericarp has been reported to contain a large amount of phenolics, including anthocyanins, procyanidins, flavonoids, lignans, and sesquiterpenes [16-18]. Dayar et al reported that concentration of anthocyanins determines the color of fruits [19]. The molecular mechanism of anthocyanin and proanthocyanidin biosynthesis in plum is inadequate. Therefore, more research is needed to reveal it.

In this study, we selected two common plum cultivars *Changli84* (Ch84) and *Dahuangganhe*(D) in Northeast China. The fruit color of Ch84 is red, while D is yellow. In order to reveal the mechanism of fruit color changes between the two cultivars, transcriptome sequencing and metabonomic analyzing were performed at three different developmental stages: young fruit stage, colour-change stage, and maturation stage. We hypothesize that anthocyanin synthesis related genes and anthocyanin metabolites play an important role in fruit color formation. This study might provide evidence for the correlation between anthocyanins and plum fruit color.

## Materials and methods

### Plant materials

All plum fruit samples were collected from the experimental farm of the Experimental farm of Jilin Academy of Agricultural Sciences, Gongzhuling City, Jilin Province. Two varieties of plum trees, Changli 84 (Ch84) and Dahuangganhe (D), were used in this study. All plum trees (six years old) grow under natural conditions, using conventional irrigation and fertilization strategies. The climate of the experimental area is temperate and monsoonal with a mean annual temperature of 5.6°C. The average annual precipitation is 594.8 mm, of which 80% falls from May to September. The fruit tissues were collected at young fruit stage (abbreviated as Y, 18^th^ May, 30 days after anthesis), colour-change stage (abbreviated as C, 17^th^ Jun, 60 days after anthesis) and maturation stage (abbreviated as D, 17^th^ July, 90 days after anthesis). Three individuals were selected for each variety, and three fruits were randomly selected from one individual for each sampling stage, and then pooled as a repeat. A total of 18 samples were collected (two varieties, three stages, three repetitions / stage / variety). For molecular analysis, tissue samples were directly snap-frozen in liquid nitrogen and kept at -80 °C.

### RNA isolation and sequencing

Total RNA extraction, library construction and RNA-Seq were performed by **Genedenovo Biotechnology** Corporation (Guangdong, China). Briefly, total RNA of these 18 samples were extracted using the TruSeq RNA Sample Preparation Kit (Autolab Biotechnology, Beijing) following the manufacture’s Guide. The quality and quantity of the purified RNA were evaluated by 1% agarose gel, and by NanoDrop ND1000 spectrophotometer (NanoDrop Technologies, Wilmington, USA). The RNA integrity number (RIN) values (>7.0) were assessed using an Agilent 2100 Bioanalyzer (Santa Clara, CA, USA). The RNA-seq libraries were prepared using an Illumina TruSeq Stranded RNA kit (Illumina, San Diego, CA). Then, the resultant 18 RNA-seq libraries were deep sequenced on an Illumina nova 6000 sequencer under the pair-end 150bp mode.

### Transcriptome data analysis

The raw reads from transcriptome sequencing were filtered by fastp (version 0.18.0) with the default parameters. Then the clean reads were assembled by Trinity (version 2.8.4) software with the default parameters. Assembled sequences Unigenes were annotated by BLASTx [20] against seven public databases in the case of >90% identity and an E-value of <0.00001, including Nr, Nt, SwissProt, KOG, Pfam, GO, and KEGG. Unigene expression levels were calculated based on FPKM values. Then, DEGs among the sample groups were identified using the DESeq2 software [21]. Differently expressed genes (DEGs) were identified based on a false discovery rate (FDR) < 0.05 and |log2 foldchange| ≥ 2. DEGs were then submitted to GO (www.geneontology.org/docs/go-enrichment-analysis/) and KEGG enrichment (www.kegg.jp/kegg/keggl.html) analysis to annotate their biological function and significantly metabolic pathways. The submission of DEGs to GO and KEGG enrichment was performed using GOseq [22] and KOBAS 2.0 [23], respectively.

### LC-MS and LC-MS/MS Analysis

In order to study the difference of metabolomics between the two cultivars at different developmental stages, untargeted metabolomics was carried out using a previously described method [24]. The freeze-dried samples were crushed using a mixer mill (MM 400, Retsch) with a zirconia bead for 1.5 min at 30 Hz. Then 100 mg powder was weighed and extracted overnight at 4°C with 1.0 ml 70% aqueous methanol containing 0.1 mg/L lidocaine for internal standard. Following centrifugation at 10000 g for 10min, the supernatant was analyzed using an LC-ESI-MS/MS system. Briefly, 2 μl of samples were injected onto a Waters ACQUITY UPLC HSS T3 C18 column (2.1 mm*100 mm, 1.8 µm) operating at 40°C and a flow rate of 0.4 mL/min. The mobile phases used were acidified water (0.04 % acetic acid) (Phase A) and acidified acetonitrile (0.04 % acetic acid) (Phase B). Compounds were separated using the following gradient: 95:5 Phase A/Phase B at 0 min; 5:95 Phase A/Phase B at 11.0 min; 5:95 Phase A/Phase B at 12.0 min; 95:5 Phase A/Phase B at 12.1 min; 95:5 Phase A/Phase B at 15.0 min. The effluent was connected to an ESI-triple quadrupole-linear ion trap (Q TRAP)–MS. The operation parameters were listed in **Table S1**.

### Metabolomic data analysis

To produce a matrix containing fewer biased and redundant data, peaks were filtered to remove the redundant signals caused by different isotopes, in-source fragmentation, K+, Na+, and NH4+ adduct, and dimerization. According to the detected peaks, metabolites were identified by searching internal database and public databases [MassBank (https://massbank.eu/MassBank/), KNApSAcK (www.knapsackfamily.com/KNApSAcK/), HMDB (https://hmdb.ca/), MoTo DB [25], and METLIN [26]). Orthogonal projection to latent structures-discriminant analysis (OPLS-DA) was performed for classification and discriminant analysis of the samples. OPLS-DA was applied in comparison groups using R software according to a previous report [27]. A variable importance in projection (VIP) score of (O)PLS model was applied to rank the metabolites that best distinguished between two groups. The threshold of VIP was set to 1. In addition, T-test was also used as a univariate analysis for screening differential metabolites. Those with a P value of T test <0.05 and VIP ≥ 1 were considered differential metabolites between two groups. Then, metabolites were mapped to KEGG metabolic pathways for pathway analysis and enrichment analysis.

## Results

### Fruit performance on two plum varieties at the developmental stage

After 90 days of growth, plums gradually matured. There were obvious differences between the two varieties. Ch84 had bigger fruit size than D. The fruit flesh color of ch84 was red and D was yellow (**Figure 1A**). The fruit weight, longitudinal diameter and transverse diameter of Ch84 were significantly higher than those of D (**Figure 1B to D**). The most striking difference is the color of the fruit.

**Figure 1.**
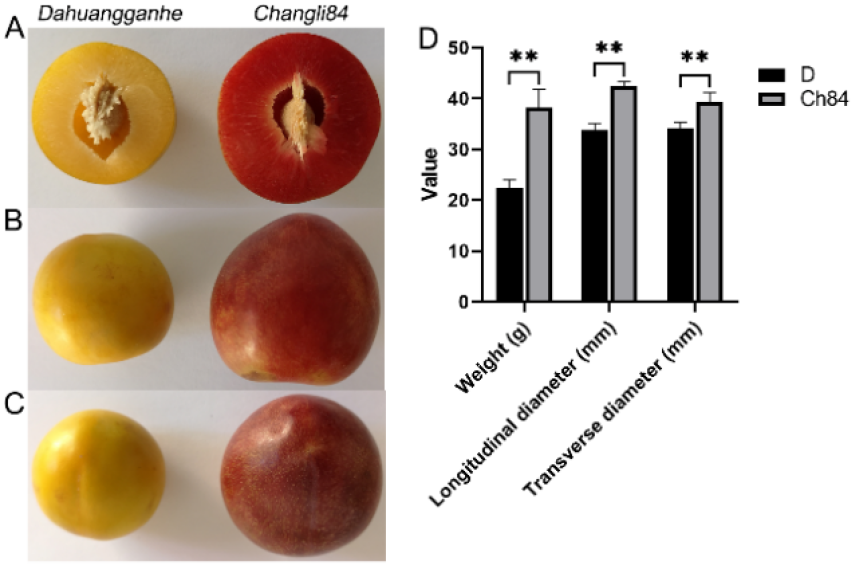
Fruit performance of the two plum varieties. All the libraries of the plum fruits produced 1,089,530,564 raw reads (150bp length / read) with an average 60,529,476 reads per library. After filtering out low quality reads, 1,087,632,936 reads (99.83% of the raw data) were obtained. The Q20 and Q30 value was 97.59% and 93.07%, respectively. The summary of sequencing data was listed in **Table S2**. The high-quality reads were assembled into 37,151 transcripts with an average length of 1216 bp and an N50 value of 2063 by Trinity software. The range of transcripts length was 201–16,179 bp. The percentages of mapping to transcript were 89.69 and 87.37%, respectively. Moreover, we also evaluated the correlation between biological repeat samples by PCA and sample to sample correlation analysis. The results showed that samples in the same group were clustered together (**Figure S1A**) and had high correlation coefficient (**Figure S1B**), indicating that the difference between biological repeat samples was small.

### Transcripts annotation

Due to lacking of a reference genome in plum, all the assembled transcripts were blasted against six public databases (Nr, Nt, SwissProt, KOG, Pfam, GO and KEGG) using search tools. Totally, 26055 transcripts were annotated using these databases. The annotation summary was shown in **Figure S2A**. There were 12,271 transcripts were commonly annotated in these databases (**Figure S2B**). The species distribution with the greatest number of plum were *Prunus mume* (60.61%), Prunus persica (8.72%), Malus domestica (3.90%), Pyrus x bretschneideri (2.92%), Populus trichocarpa (2.86%), etc (**Figure S2C**).

### Analysis of differentially expressed genes (DEGs) between the two plum varieties

According to false discovery rate (FDR) < 0.05 and |log2 foldchange|≥ 2, DEGs were identified using the DESeq software package. As shown in **Figure 2A**, 4,994, 6,696 and 5,322 DEGs were identified in comparison of Ch84-Y *vs*. D-Y, Ch84-C *vs*. D-C, and Ch84-D *vs*. D-D, respectively. The venn analysis showed that 2,429 DEGs were commonly expressed in the three comparisons (**Figure 2B**). Go enrichment was performed on all the DEGs to cluster the genes with similar functions. The GO enrichment results showed that no significant BP terms enriched in Ch84-Y *vs*. D-Y and Ch84-D *vs*. D-D (**Figure 3A**). For KEGG enrichment analysis, we found that the “Phenylpropanoid biosynthesis” (ko00940), “Flavonoid biosynthesis” (ko00941), “Plant-pathogen interaction” (ko04626) and “Biosynthesis of secondary metabolites” (ko01110) were significantly enriched (**Figure 3B**). DEGs in “Biosynthesis of secondary metabolites” and “Flavonoid biosynthesis” were the same, including *PKS5, CHS, ACT, CHI3, FLS, CYP, HCT2, DFR* etc. DEGs in “Phenylpropanoid biosynthesis” included *CHS, CAD1, 4CL2, ACT, GT5, PER*, etc. Also, RPM1, CNGC1, XA21, RPP13, Hsp90ab1, MYB62, etc were significantly enriched in “Plant-pathogen interaction”. In these pathways, “Flavonoid biosynthesis” is closely related to anthocyanins, and the expression levels of related genes are shown in **Figure 3C**.

**Figure 2.**
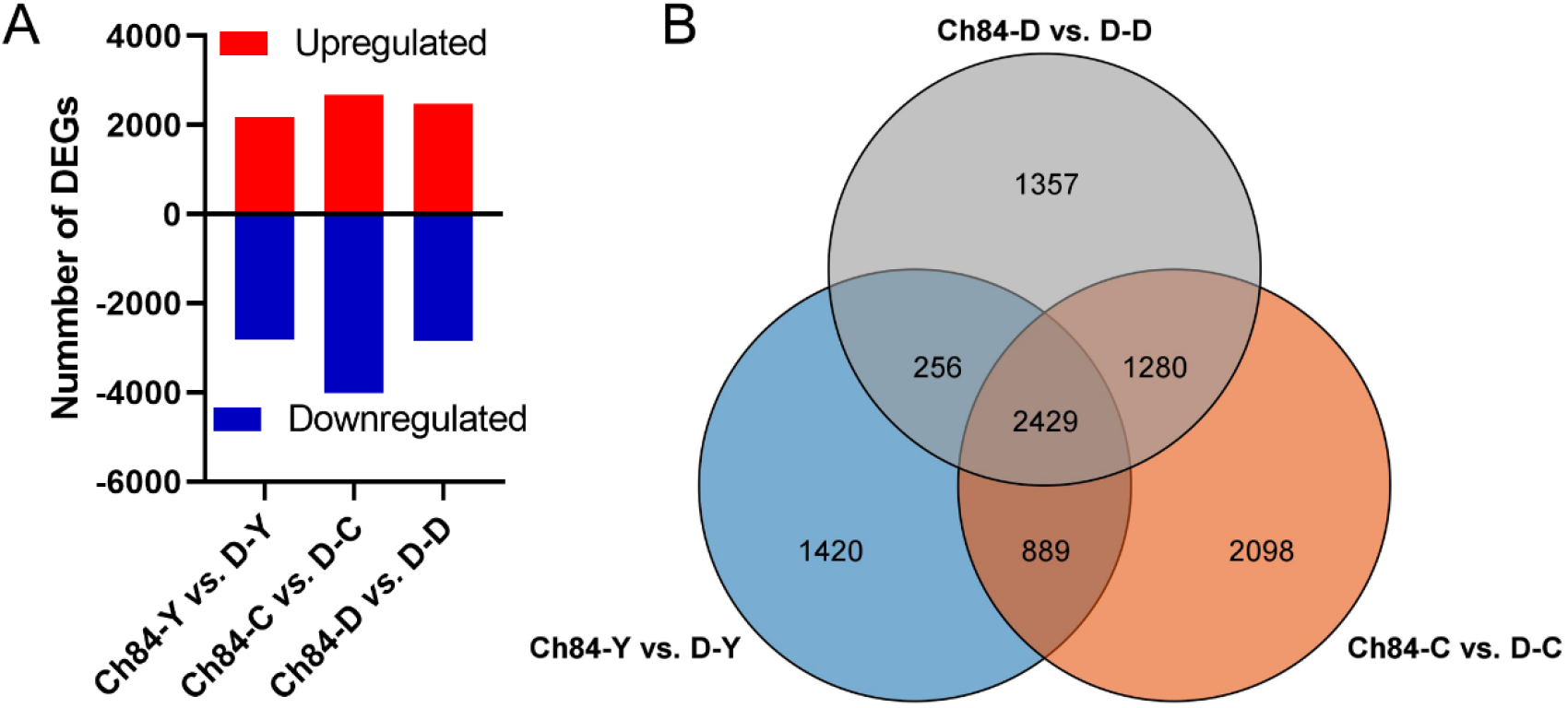
Statistics of differentially expressed genes (DEGs) A shows the number of DEGs between the two plum varieties at different developmental stage. B shows the Venn analysis results of different comparisons.

**Figure 3.**
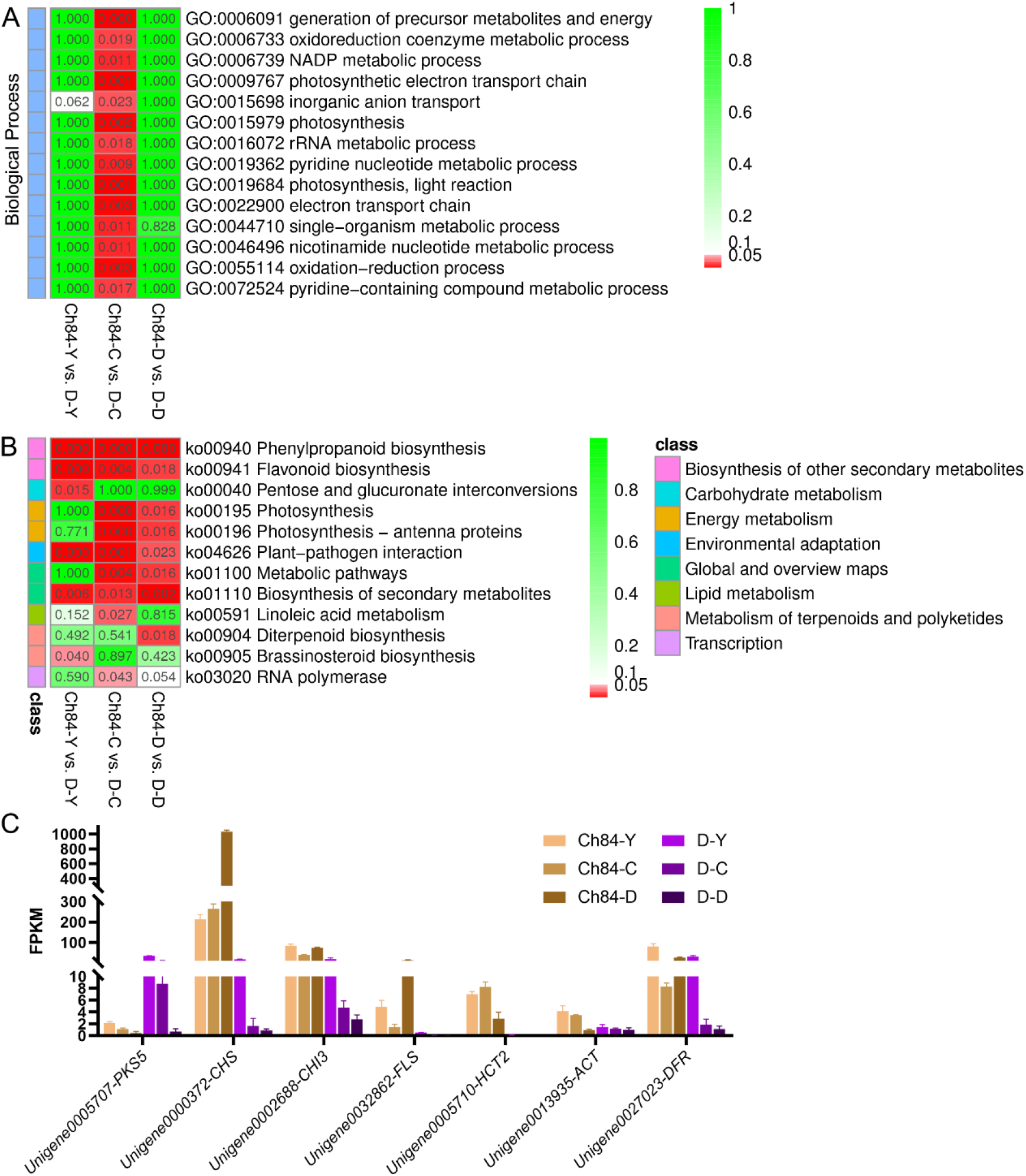
The GO and KEGG enrichment results of DEGs and the expression of candidate DEGs. A and B shows the GO and KEGG enrichment results of all the DEGs, respectivly. The redder the color, the higher the significance. C shows the expression of candidate DEGs of the two plum at different developmental stage.

### Transcriptional control of MBW (MYB–bHLH–WD) protein complexes

In order to further explore the DEGs that related to fruit color, we identified the gene expression level of MBW protein complexes. We found that 6 *MYB* related DEGs were highly expressed in Ch84 but lowly in D. Nine *bHLHs* had a significantly higher expression in Ch84 than D. However, there was no *WD40* differently expressed between Ch84 and D. By mapping the gene expression heat map of MBW complex, *MYB* and *bHLH* may be the key genes regulating anthocyanin synthesis in plum (**Figure 4**).

**Figure 4.**
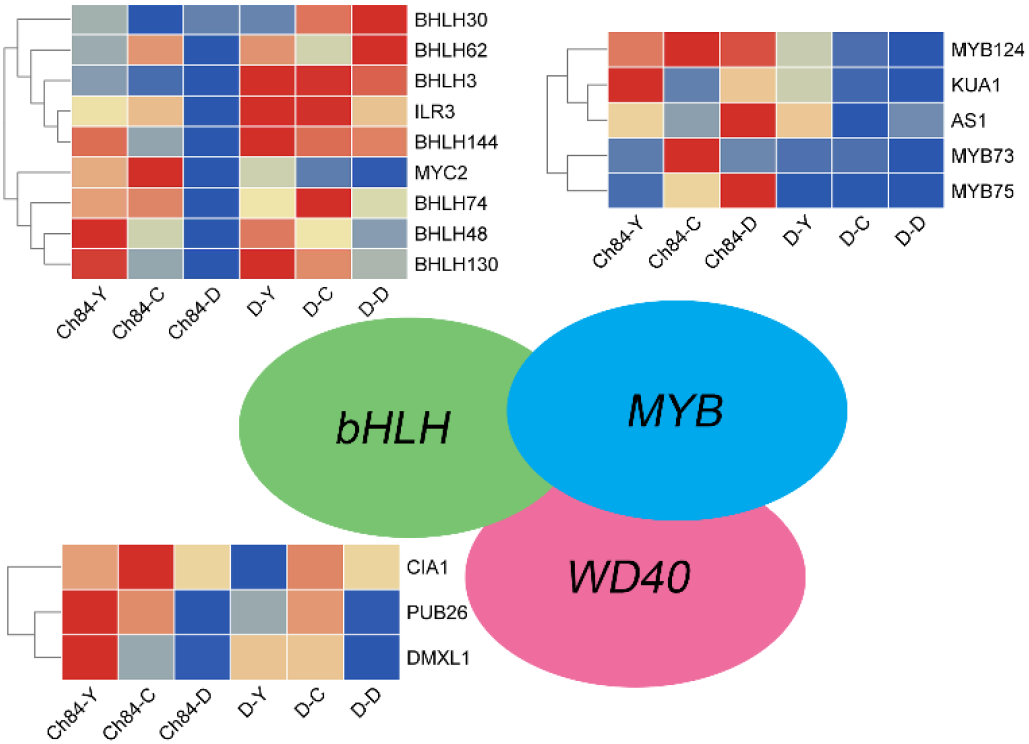
Expression of MBW protein complexes. A simplified model depicting the transcription factors of MYB, bHLH and WD40. The redder the color of the box, the higher the expression, and the bluer the color, the lower the expression

### Analysis of differential metabolites (DMs) between the two plum varieties

To distinguish the classes and to assess the global metabolism variations, the unsupervised PCA was performed in both positive and negative spectra. The PCA score plot of positive spectra indicated that there was a clear classification of observations of Ch84 and D (**Figure 5A**). To further distinguish the Ch84 and D and to identify differential variants, a supervised OPLS-DA was conducted. In **Figure 5B, C and D**, a remarkable separation of LC-MS data in Ch84 and D groups at different developmental stages was observed in the OPLS-DA score plot, indicating that this OPLS-DA model was non-overfitting. The results of these assessments mentioned above suggested that the LC-MS data quality was reliable. Fifty-four differential metabolites were identified and the heatmap of the differential metabolites were shown in **Figure 6A**. To uncover the most relevant biological pathways of osimertinib resistance, the KEGG enrichment analysis was performed. The enrichment and topology analysis demonstrated that main difference of metabolites between the two varieties was related to “Cysteine and methionine metabolism” (ko00270), “Biosynthesis of amino acids” (ko01230) and “Aminoacyl-tRNA biosynthesis” (ko00970) (**Figure 6B**). Interestingly, we found significant differences in the content of several anthocyanin related metabolites between the two cultivars, including Cyanidin 3-glucoside, Cyanidin-3-O-alpha-arabinopyranoside, Procyanidin B2 and Procyanidin B1. The level of these mentioned metabolites were shown in **Figure 6C**.

**Figure 5.**
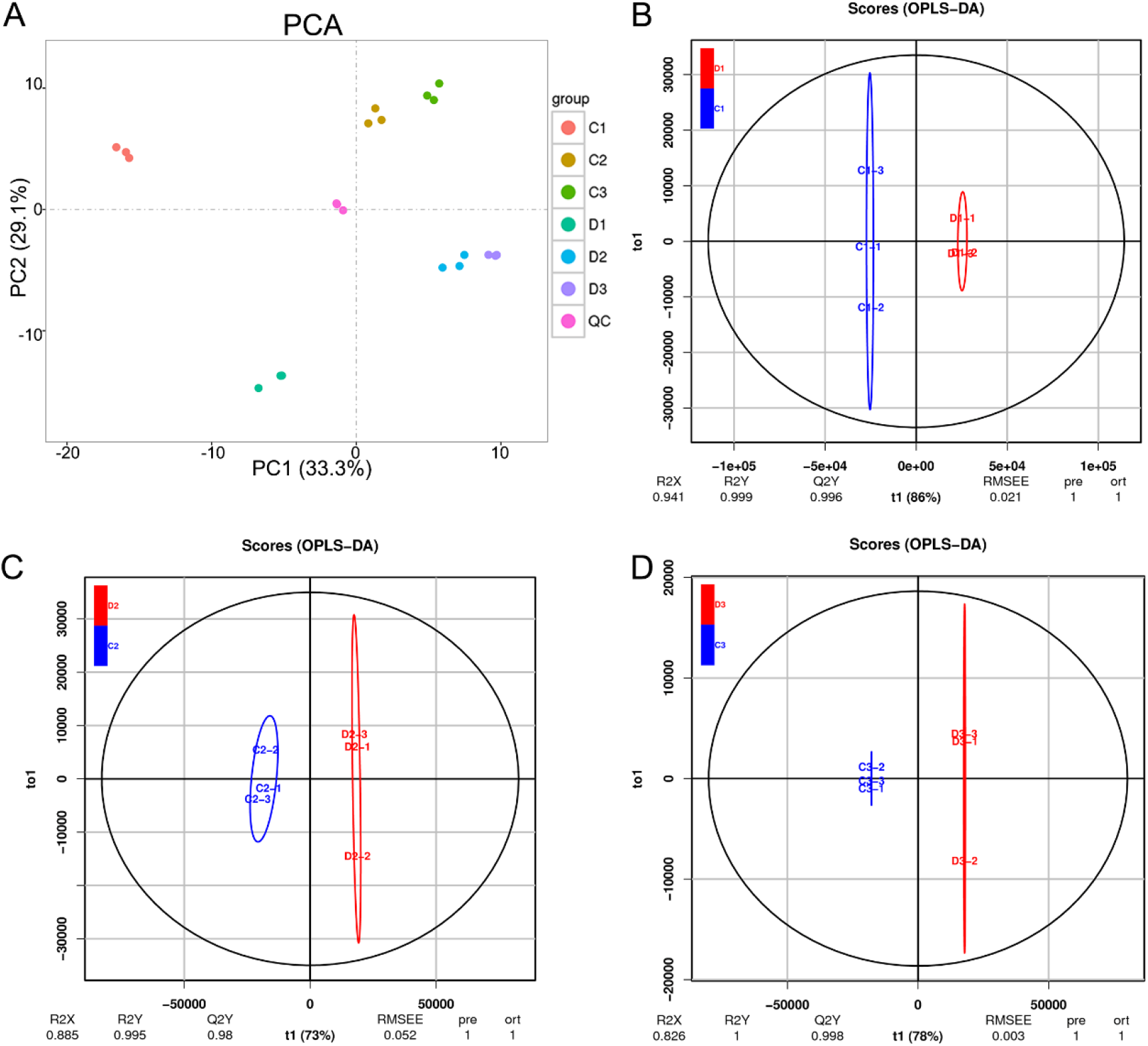
Summary of the metabolomic data. A shows the PCA results, different colors represent different groups. B to D shows the OPLS-DA analysis results between the two plum varieties at the young fruit stage, color changing stage and fruit mature stage.

**Figure 6.**
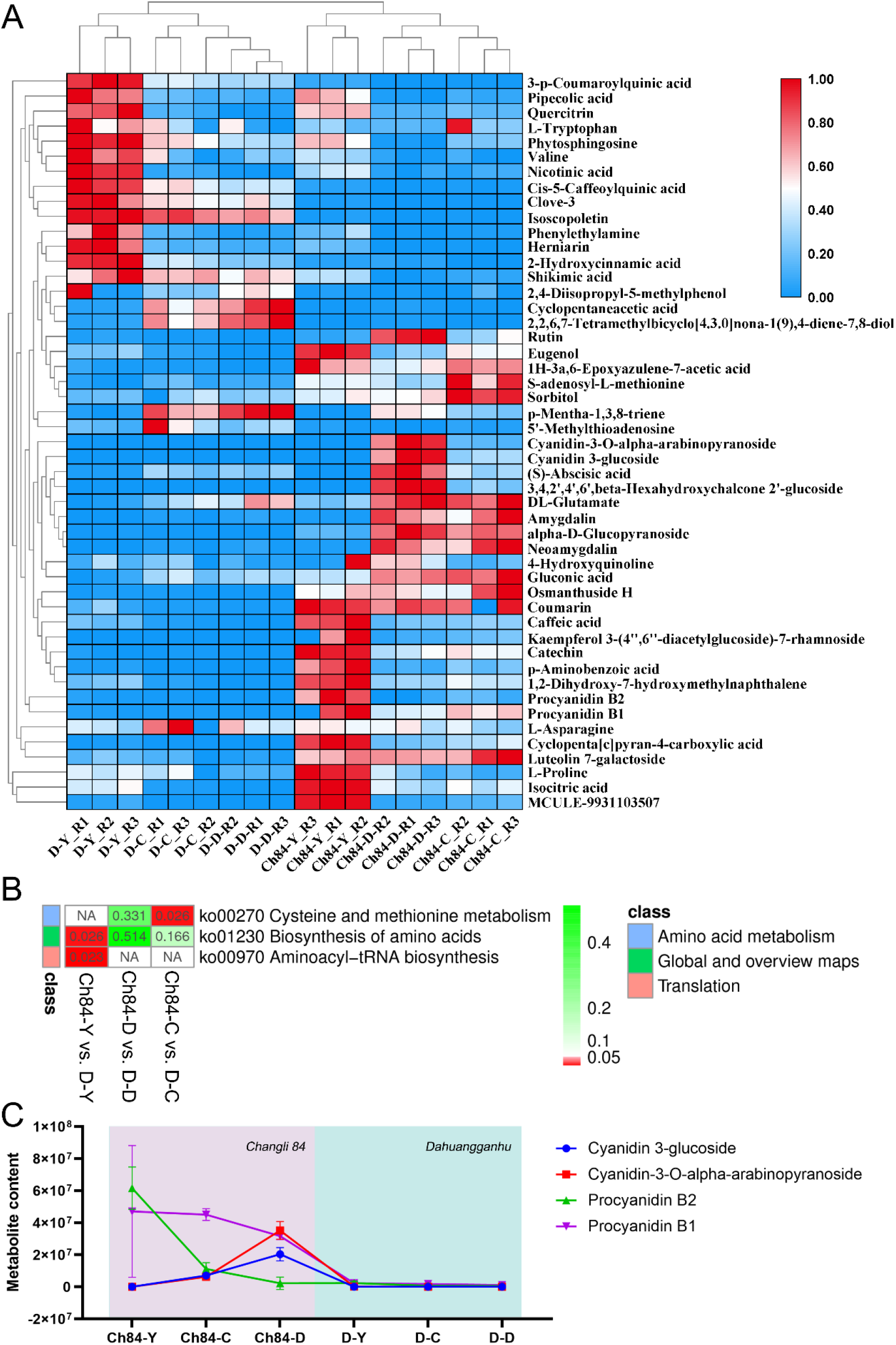
Differential metabolites and functional enrichment analysis. A shows the level of differential metabolites. The redder the color, the higher the level. B shows the KEGG enrichment results based on these differential metabolites. C shows the level of candidate differential metabolites of the two plum at different developmental stage.

## Discussion

There are obvious differences in fruit color between the two plum varieties. Finding out the difference can help us understand the mechanism of fruit color. As shown in **Figure 1**, The fruit color of Ch84 is dark red, while that of D is yellow. The main reason for this difference may be caused by different anthocyanin content in the plum fruit. The transcriptome analysis results showed that “ Flavonoid biosynthesis” was significantly enriched in the KEGG analysis. Flavonoid biosynthesis is one of the most extensively studied secondary metabolic pathways in plants [28, 29]. Flavonoid biosynthesis is regulated by a complex network of signals triggered by internal metabolic cues and external signals, including visible light, pathogen attack, nitrogen, phosphorus, and iron deficiencies, temperature, etc [30]. Genetic characteristics are also a factor that cannot be ignored. The results of flavonoid biosynthesis related gene expression showed that their expression in the two plum cultivars were quite different. In particular, the expression trend of some DEGs was opposite between the two cultivars, such as *CHS, DFR* and *FLS*. Anthocyanin synthesis is a process that involves many steps [31-33]. *PAL, CHI, CHS, F3H, DFR, ANS* and *UFGT* are closely related with anthocyanin biosynthesis [29, 34]. *CHS* is responsible for initiating flavonoid biosynthesis [35] and F3GT has been shown to be responsible for anthocyanin biosynthesis [36]. Peng reported that *CHS* and *F3GT* are crucial for total anthocyanin accumulation [37]. In this present study, the expression of *CHS* increased with the growth of fruit in Ch84, but decreased in D, indicating that the anthocyanins in Ch84 were accumulated continuously, while those in D were consumed. Correspondingly, the results of LC-MS data also confirmed that anthocyanin content changed from procyanidin to cyanidin. In addition, *CHI* maintained a high level in Ch84, but decreased in D. In general, the flavonoid pathway begins with the sequential condensation of one molecule of 4-coumoryl-CoA and three molecules of malonylCoA by *CHS*, resulting in the formation of naringenin chalcones [38]. The chalcone is then stereospecifically isomerised to the flavanone naringenin by the enzyme CHI [39]. Hence, the expression of *CHI* also plays an important role in the accumulation of anthocyanin. According to our results, the activity of flavonoid biosynthesis is determined by varieties, and the high expression of key genes leads to the changes of fruit color.

It is usually considered that anthocyanin biosynthesis is regulated by MBW complexes consisting of different MYBs, bHLHs and WD40 transcription factors [40]. But there are exceptions. Myb and bHLH transcription factors can also co-regulate the expression of anthocyanin biosynthesis genes without depending on WD40 protein [41]. The results of this study suggested that the differential expression of *MYB* and *bHLH* might be another key factor leading to fruit color change. Higher *MYBs* and lower *bHLHs* might make the fruit more red. On the contrary, the fruit with lower *MYBs* and higher *bHLHs* tends to be yellow. The mechanism needs more experiments to verify, but at least based on the transcriptome results, we believe that the expression levels of *MYBs* and *bHLHs* dominate the color change of plum fruit. The role of WD40 is limited.

The results of metabolomics also give us a lot of hints about the mechanism of plum fruit color. Polyphenols including yellow flavonoids, procyanidins (B1 and B2) and cyanidin-3-O-glucoside in substantial amounts have been characterized in different palm fruits [42]. In our results, procyanidin B1 and B2 had the highest level at young fruit stage in Ch84 and the content of procyanidin B2 decreased sharply at the color change stage. Conversely, the content of cyanidin increased with the growth of fruit and reached the peak at the maturation stage. While for D, the metabolites mentioned above did not change significantly at all developmental stages. As we know, procyanidins are members of the proanthocyanidin (or condensed tannins) class of flavonoids. They are oligomeric compounds, formed from catechin and epicatechin molecules. They yield cyanidin when depolymerized under oxidative conditions [43]. Therefore, we speculated that the content of polyphenols, like procyanidin B1 and B2, in plums at young fruit stage might be the leading factors of the matured fruit color.

## Conclusion

Anthocyanins content is the main factor for the difference of fruit color between the two varieties. The content of procyanidin B1 and B2 in plums at young fruit stage might be the leading factors of the matured fruit color. At the maturation stage, the cyanidin produced by procyanidins keeps the color of the fruit red. Correspondingly, genes in “flavonoid biosynthesis” pathway are active. DEGs like *CHS, CHI3, DFR* had higher expression level in Ch84 than those in D, and these genes regulated the accumulation of anthocyanin in plum. Also, we speculated that the expression levels of *MYBs* and *bHLHs* might dominate the color change of plum fruit, while the role of WD40 is limited.

## Data Availability Statement

The transcriptional datasets analyzed during the current study are available in the National Center for Biotechnology information (NCBI) repository, and the BioProject accession number was PRJNA726302(https://www.ncbi.nlm.nih.gov/bioproject/?term=PRJNA726302). The Prunus salicina transcriptome shotgun assembly (TSA) project has the project accession GJEK00000000.1. This version of the project (01) has the accession number GJEK00000000.1, and consists of sequences GJEK01000001-GJEK01035577.

